# Developmental Chlorpyrifos Exposure Disrupts Tectal Circuit Formation and Behavior in *Xenopus laevis*

**DOI:** 10.64898/2026.05.29.728774

**Authors:** Virgilio Lopez, Alyssa Rust, Adrian C Thompson, Zain Peerbhoy, Carlos D Aizenman

**Affiliations:** Brown University, Department of Neuroscience; Boston University, Department of Neuroscience

## Abstract

Developmental exposure to organophosphate pesticides has been associated with adverse neurodevelopmental outcomes, but the circuit-level mechanisms underlying these effects remain poorly understood. Here, we examined how low-level chlorpyrifos (CPF) exposure affects neural circuit maturation and behavior in *Xenopus laevis* tadpoles. Tadpoles were exposed to 1 µM CPF from developmental stage 42 to stage 49, spanning a critical period of synaptogenesis and circuit refinement. CPF-exposed tadpoles displayed abnormal schooling behavior, characterized primarily by impaired body-axis alignment despite preserved group aggregation, as well as altered spontaneous swimming marked by reduced looping and increased seizure-like activity. Whole-cell recordings from tectal neurons revealed persistent reductions in inhibitory synaptic drive, consistent with altered excitation–inhibition balance. Although acute CPF exposure transiently increased intrinsic excitability of tectal neurons as well as baseline swimming activity, this effect was not maintained after chronic exposure. Morphological analysis of GFP-labeled tectal neurons revealed altered dendritic branching distribution despite no change in total dendritic length or branch number. Together, these findings suggest that developmental CPF exposure disrupts tectal circuit maturation, leading to abnormal neural connectivity and maladaptive behavioral outcomes relevant to neurodevelopmental dysfunction.

## Introduction

Neurodevelopmental disorders such as autism spectrum disorder (ASD) are primarily genetic in origin, with heritability estimates as high as 90% (Sandin et al., 2017). However, these conditions are also highly heterogeneous, both in symptomatology and developmental trajectory. This variability suggests that additional non-genetic factors, particularly those acting during critical periods of brain development, play a crucial role in shaping risk and phenotype. This highlights the need to understand how environmental exposures, particularly those occurring pre and perinatally, can alter the trajectory of neurodevelopment. Among such exposures, agricultural chemicals are of particular concern given their ubiquity and documented capacity to cross the placenta during sensitive developmental windows.

Organophosphate pesticides (OPs) are among the most widely used chemical agents in the agricultural industry. As of 2021, approximately 40% of pesticides used in agriculture were OPs, and these function through irreversible inhibition of acetylcholinesterase (AChE), which can result in adverse health outcomes (Kaushal et al., 2021). One of the most widely used OPs is Chlorpyrifos (CPF). This compound has garnered substantial attention due to its complicated regulatory history in agricultural pest control. While its residential use was banned in 2000 because of health concerns, its agricultural application has undergone repeated regulatory reassessment, where a complete ban was proposed in 2015 due to concerns about negative neurodevelopmental outcomes due to prolonged low-level exposure during prenatal or early-life periods. After a change in leadership at the EPA, this proposed ban was reversed in 2017, due to the high reliance on CPF in agricultural production, and questions about the evidence for negative neurodevelopmental outcomes (Rauh et al., 2006; US EPA, 2014; Volcovici, 2019).

Occupational exposure to concentrated CPF has been associated with acute toxicity, including respiratory failure, tachycardia, kidney injury, and seizures (H.-F. Liu et al., 2020). Mechanistically, CPF inhibits acetylcholine esterase (AChE), the enzyme responsible for breaking down synaptically-released acetylcholine (ACh), resulting in excessive cholinergic signaling and subsequent neurotoxicity (Lan et al., 2017; Silva, 2020). Although the toxicological profile of CPF in adults is well characterized, its effects on fetal neurodevelopment remain comparatively less understood. Emerging evidence links prenatal CPF exposure to developmental impairments and elevated risk of autism spectrum disorder (ASD) or other negative neurodevelopmental outcomes (González-Alzaga et al., 2014; Shelton et al., 2014; von Ehrenstein et al., 2019). CPF is highly lipophilic and capable of crossing the placenta, facilitating in utero exposure (Rauh et al., 2006). Epidemiological studies in inner-city cohorts have correlated elevated cord blood CPF levels with increased incidence of neurodevelopmental delays. Further studies showed increased incidence of ADHD, working memory deficits, and cognitive and motor impairments following prenatal exposure (Bouchard et al., 2010; Burke et al., 2017).

ASD pathology is increasingly attributed to fundamental abnormalities in neural circuit development (Pratt & Khakhalin, 2013), and CPF-induced ASD-like phenotypes linked to neural circuit development have been demonstrated across multiple model systems. In infant rats, CPF exposure altered ultrasonic vocalization patterns in a manner comparable to prenatal valproic acid (VPA) exposure, a well-established inducer of social behavioral deficits (Morales-Navas et al., 2020). In mice, CPF exposure has been associated with reduced dendritic complexity and increased prevalence of immature dendritic spines within the hippocampus (Mullen et al., 2016). In zebrafish, CPF disrupts axonal growth and neuronal connectivity and reduces dopamine levels, both neurobiological features implicated in ASD pathophysiology (DiCarlo & Wallace, 2022; Eddins et al., 2010; Yang et al., 2011).

Due to their well-characterized neurodevelopmental trajectory and experimental accessibility, Xenopus laevis tadpoles represent a powerful model for studying neural circuit development and neurodevelopmental disorders, (Gore et al., 2021; James et al., 2015; Pratt & Khakhalin, 2013). In Xenopus laevis tadpoles, developmental CPF exposure induces gross morphological abnormalities of craniofacial structures, and downregulates several neurodevelopmental genes such as fgf8, Sox9, and Bmp4 (Tussellino et al., 2016). Additional studies report altered thyroid hormone axis signaling, aberrant brain morphology, and reduced axon myelination following CPF exposure (Spirhanzlova et al., 2023). A critical gap in the current literature lies in understanding how CPF-induced anatomical alterations at the neural circuit level translate into functional behavioral outcomes that are consistent with neurodevelopmental disorders. Furthermore, the effects of low-level CPF exposure on the electrophysiological properties of developing neurons remain poorly defined.

In this study, we aim to investigate the impact of CPF exposure on the electrophysiology, dendritic architecture, and behavior of Xenopus laevis tadpoles during a critical period of synaptogenesis and circuit development in the optic tectum. Whole-cell electrophysiology will be used to assess development of intrinsic excitability and synaptic transmission. Dendritic architecture and arbor complexity will be assessed in GFP-expressing tectal neurons *in vivo*.

These data will be integrated with behavioral assays assessing motility and social behavior to determine whether CPF-induced cellular alterations correspond to maladaptive functional outcomes. This work aims to clarify the neuronal-level mechanisms underlying the neurodevelopmental consequences of CPF exposure and contribute to understanding its broader implications for ASD-related pathophysiology.

## Methods

### Rearing and CPF Exposure

All animal experiments were performed in accordance with and approved by Brown University Institutional Animal Care and Use Committee standards. Wild-type and albino Xenopus laevis tadpoles were bred in-house, and were kept in 10% Steinberg’s rearing media until they reached developmental Stage 42 (typically 5-7 days dpf; Nieuwkoop & Faber, 1994). Tadpoles were reared in an incubator held at 19-20°C with a 12:12 hour light/dark cycle. They were then transferred into either a container with either 200 ml of Steinberg’s rearing media or rearing media with 200 ml of 1 μM CPF. Tadpoles were kept in the incubator for 8-10 days and allowed to develop until stage 49. CPF stocks were prepared with methanol and methanol alone was used as a control vehicle.

### Schooling behavior

Schooling assays were carried out as previously described (Lopez et al., 2021), using the publicly available analysis code (https://github.com/khakhalin/Xenopus-Behavior). For each trial, 30 control or CPF-exposed tadpoles were placed in a 17-cm diameter glass bowl positioned on an LED tracing tablet (Picture/Perfect light pad), which rested on a dental vibrator (Jintai). Images were collected with a GoPro Hero 7 (GoPro Inc.) every 5 min for 1 h, with tadpoles dispersed by activating the vibrator 150 s prior to each frame. For analysis, tadpole position and heading were determined by marking x–y coordinates of the head and gut for each animal using the multipoint tool in FIJI. A Delaunay triangulation was then applied to compute inter-tadpole distances and relative angles between neighbors, and distributions were compared using a Kolmogorov–Smirnov test.

### Swimming

To quantify swimming behaviors, stage 49 tadpoles were transferred into individual wells of a 6 well plate (Corning) containing 4ml of 10% Steinberg’s media. The plate was illuminated from below with a LED tracing pad (Picture/Perfect) and imaged at 30 frames/s from above with a SCB 2001 color camera (Samsung) for 10 minutes. The center-point of each tadpole was tracked in real time using EthoVision XT (Noldus Information Technology). Tadpole motility and behavior was then quantified offline using a custom MATLAB script. Briefly, swimming bouts were detected automatically as periods of sustained motility above threshold (>0.2cm/s), with the type of behavior being performed defined as periods of looping, darting, turning, or by irregular movements such as seizing or spiraling.

### Electrophysiology

For whole-cell recordings from tectal neurons, tadpole brains were prepared as previously described to expose the ventral surface of the tectum (Ciarleglio et al., 2015; Thompson & Aizenman, 2024; Wu et al., 1996). Briefly, tadpoles were anesthetized in 0.02% tricaine methanesulfonate (MS-222), after which brains were opened along the dorsal midline, removed, and pinned to a sylgard block in a recording chamber at room temperature. Brains were maintained throughout the experiment (typically 2–3 h) in HEPES-buffered extracellular saline (in mM: 115 NaCl, 2 KCl, 3 CaCl₂, 3 MgCl₂, 5 HEPES, 10 glucose; pH 7.2; osmolarity 250 mOsm). The ventricular membrane was removed by gentle suction with a broken glass pipette to access principal tectal neurons. Recordings were limited to the middle third of the tectum to minimize developmental variability along the rostrocaudal axis (Khakhalin & Aizenman, 2012; Hamodi & Pratt, 2014). Cells were visualized using a Nikon Eclipse E600FN microscope with a 60× water-immersion objective, and whole-cell recordings were obtained using 10–12 MΩ glass micropipettes. Signals were amplified with a Multiclamp 700B, digitized at 10 kHz with a Digidata 7550B, and acquired in pClamp 11 (Molecular Devices). Active currents were isolated by real-time leak subtraction. Membrane potential was not corrected for the predicted 12 mV liquid junction potential. Neurons with series resistance >50 MΩ were excluded. Data were analyzed using Axograph X (John Clements).

Recording pipettes were filled with K-gluconate–based intracellular saline (in mM: 100 K-gluconate, 8 KCl, 5 NaCl, 1.5 MgCl₂, 20 HEPES, 10 EGTA, 2 ATP disodium salt hydrate, 0.3 GTP sodium salt hydrate; pH 7.2; osmolarity 255 mOsm). For spontaneous EPSC and IPSC recordings, cells were voltage-clamped at −45 mV and +5 mV, respectively, corresponding to the reversal potentials for inhibitory and excitatory currents in the tectum. Spontaneous synaptic events were detected using a template-matching algorithm in Axograph X.

### Electroporation and Neuronal Morphology

To label single tectal neurons, albino tadpoles were anesthetized with 0.02% MS-222 and then whole-brain electroporation was used to transfect tectal neurons with pCALNL-GFP plus pCAG-Cre plasmids (Schohl et al., 2020). Two platinum electrodes between 1-2mm were placed on the skin overlaying both sides of the tadpole midbrain. Using a glass micropipette, the plasmids were injected into the middle ventricle of the brain. After the injection, between three to four electrical pulses at 50V with an exponential decay of 70 ms were delivered to incorporate the DNA into the tectal cells. The tadpoles were then returned to their respective media to recover and allowed to develop until stage 49. Electroporated tadpoles were screened with a fluorescence microscope to determine which animals expressed the plasmid in the tectal region. Tadpoles required at least one tectal neuron showing clear GFP expression to be included for imaging.

A Zeiss LSM 800 confocal microscope was used to image GFP expressing neurons in vivo in anesthetized tadpoles. The 3D morphology of transfected neurons was reconstructed using the open-source software Neutube (https://www.neutracing.com/). The 3D reconstructions were then analyzed using Fiji (https://imagej.net/software/fiji/) to find the total dendritic length, average dendritic length, and number of dendritic branches. A Sholl analysis was also conducted in Fiji to analyze dendritic branching patterns.

### Analysis and Statistics

For all experiments groups were blinded to the experimenter where possible. When blinding was not possible during experimental acquisition data was analyzed blind to experimental conditions. For schooling experiments each experimental N represented one experimental run, and for all experiments at least three different clutches of tadpoles were used. Non-parametric statistics were used where appropriate. Further details are included in each experimental or results section.

## Results

Between developmental stages 42 and 49, Xenopus laevis tadpoles go through a critical developmental time period in which neurons in the optic tectum undergo significant maturation in their intrinsic, synaptic, dendritic and circuit level properties (Akerman & Cline, 2006; Pratt & Aizenman, 2007). This correlates with refinement of multisensory inputs to the tectum and the maturation of sensory guided behavior (Deeg et al., 2009; Dong et al., 2009; Felch et al., 2016; Pratt & Aizenman, 2009). To determine whether chronic exposure to CPF during this developmental window leads to abnormal maturation of tectal circuitry, we first examined the effects of CPF exposure on schooling behavior, which emerges by stage 49 and depends on the normal development of multisensory circuits as well as ability to respond to social cues from conspecifics (James et al., 2015). To determine a working concentration of CPF we referred to the literature, which showed that exposure levels above 0.5 mg/L (∼1.5 µM) resulted in morphological abnormalities and death of Xenopus tadpoles (San Segundo et al., 2013). This was confirmed with a series of experiments that showed that developmental exposure between stages 42 and 49 to 1 µM CPF resulted in full survival and no gross abnormalities in development, while higher concentrations proved to be detrimental (**Table 1**). Thus, for all developmental exposures, tadpoles were reared in standard media until developmental stage 42 when they were introduced to rearing media containing 1 µM CPF (typically 6-7 dpf) and allowed to develop until stage 49 (typically 14-16 dpf).

**Table 1.** Survival of tadpoles exposed to different concentrations of CPF between Stages 42 and 49. Also indicated percent of surviving tadpoles with swimming abnormalities and percent of total tadpoles with severe physical deformities.

| Treatment (CPF μM) | Control | Vehicle | 0.1 | 0.5 | 1 | 5 |
| --- | --- | --- | --- | --- | --- | --- |
| % Dead | 0 | 0 | 0 | 0 | 0 | 100 |
| % Abnormal swimming | 0 | 0 | 0 | 32 | 72 | 0 |
| % Morphological Abnormality | 0 | 0 | 0 | 0 | 0 | 100 |
| N | 50 | 50 | 25 | 25 | 50 | 25 |

### Schooling

In the absence of environmental stressors, tadpoles exhibit social behavior in the form of schooling when put into a large arena. Schooling is quantified by comparing inter-tadpole distances between nearest neighbors, as well as intertadpole angles (Lopez et al., 2021). We compared the schooling behavior of control and tadpoles reared in 1 µM CPF (chronic CPF). We found small but significant differences in the distributions of inter-tadpole distances between the two groups of tadpoles (**Fig. 1A**, **p < 0.001 K-S, *N* = 59 and 56 tests for control and Chronic 1µM CPF, respectively**), with the control treated group indicating slightly tighter clustering, yet indicating that CPF exposed tadpoles still aggregate into groups. In contrast, body-axis orientation (as measured by inter-tadpole angles) differed significantly between conditions. Small inter-tadpole angles indicate that tadpoles are swimming in a similar direction as their neighbors as they school. CPF-reared tadpoles exhibited abnormal co-orientation within schooling clusters, with a more random distribution of inter-tadpole angles (Fig 1B **p < 0.01 K-S, *N* = 59 and 56 tests for control and Chronic 1µM CPF, respectively**). Together, these results show that while CPF-reared tadpoles were able to cluster into groups, their abnormal body-axis orientations indicate abnormal schooling behavior (Fig. 1C).

**Figure 1.**
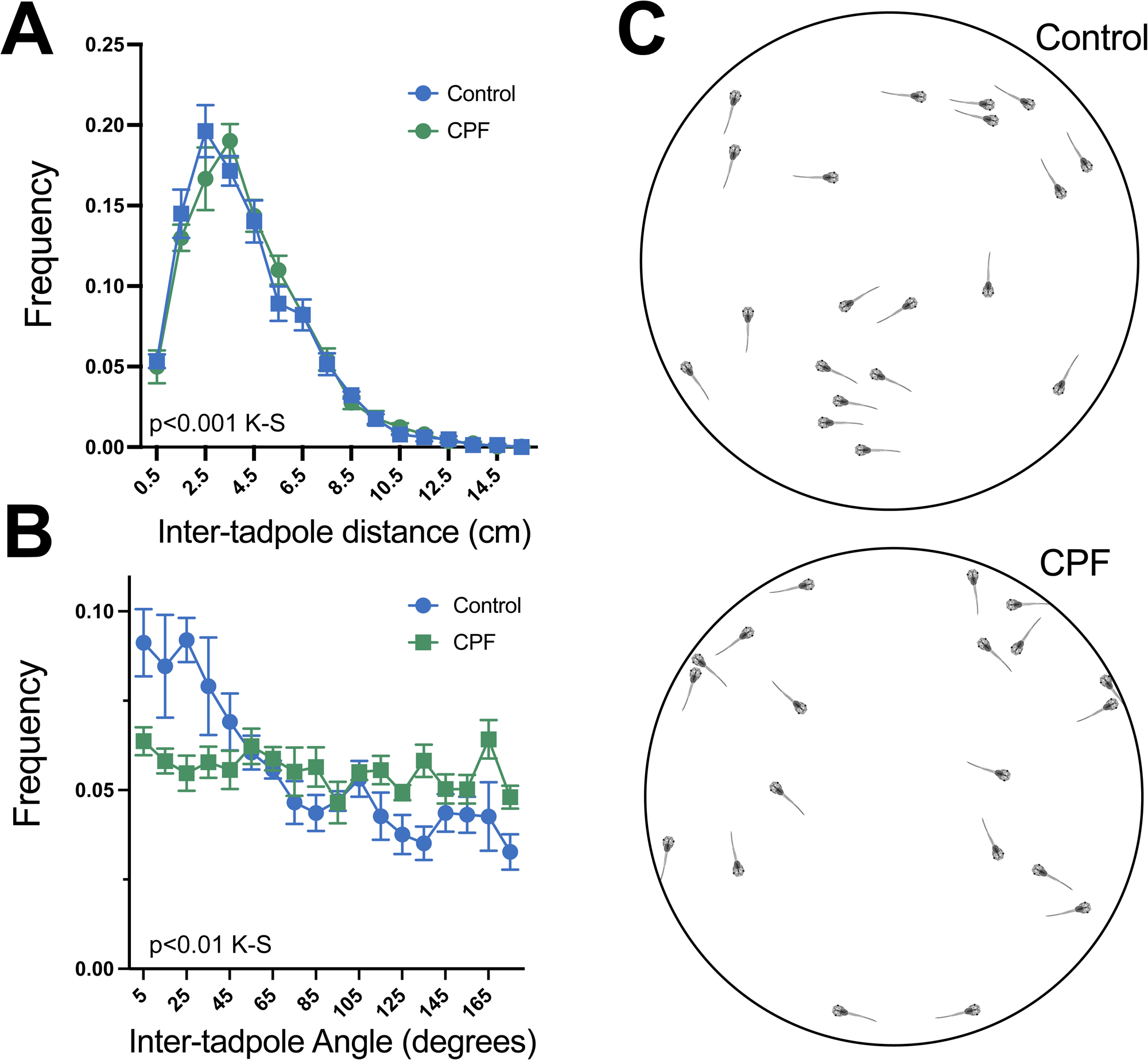
Developmental CPF exposure disrupts schooling behavior. **(A)** Frequency distributions of inter-tadpole distances between nearest neighbors for control (blue) and chronically CPF-exposed tadpoles (1 µM CPF from stage 42 to stage 49; green). Both groups show similar peaked distributions indicating that tadpoles aggregate into groups under both conditions, although the distributions differ slightly, with control tadpoles showing somewhat tighter clustering. **(B)** Frequency distributions of inter-tadpole angles, where small angles indicate that neighboring tadpoles are oriented in similar directions. Control tadpoles show a distribution skewed toward small angles, consistent with co-oriented schooling, whereas CPF-exposed tadpoles show a more uniform distribution across angles, indicating loss of body-axis alignment. Data are shown as mean ± SEM across runs. **(C)** Representative tracings from single frames of tadpole positions and headings within the schooling arena for control (top) and CPF-exposed (bottom) groups. Control tadpoles tend to align with neighbors, whereas CPF-exposed tadpoles cluster into groups but with more randomly oriented body axes.

### Swimming behavior

One possibility is that the abnormal schooling behavior may result from differences in baseline locomotor activity, such as an inability to swim normally. To determine if CPF exposure affects baseline locomotor activity, we analyzed the swimming behavior of tadpoles exposed to 1 µM CPF during development, and following acute exposure for 1 hour. Tadpoles were then tracked for 10 minutes in individual wells to measure spontaneous swimming (Fig. 2A-C).

**Figure 2.**
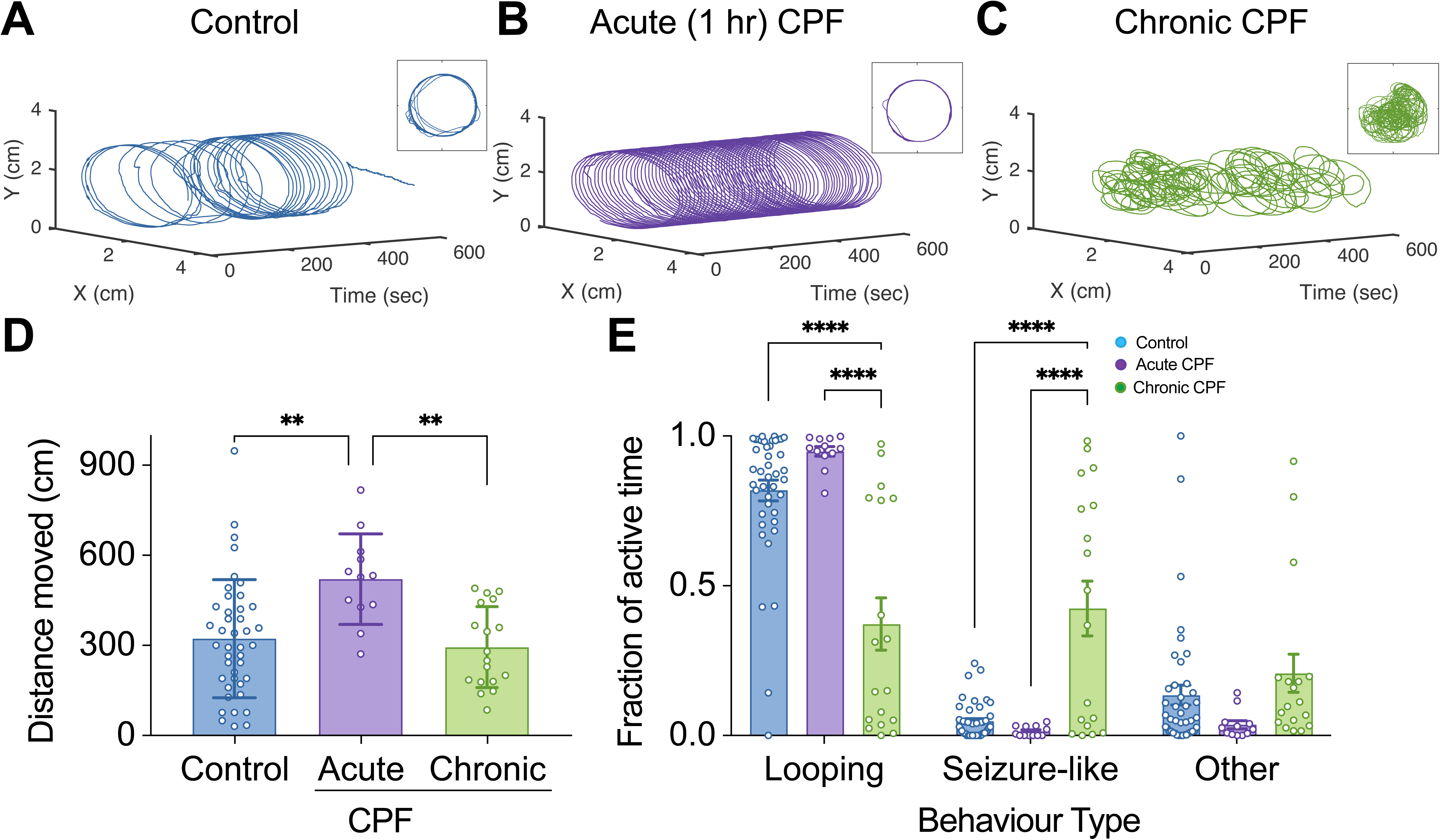
Acute and chronic CPF exposure differentially affect spontaneous swimming behavior. **(A–C)** Representative 10-minute swimming trajectories for individual tadpoles from each condition, plotted as X–Y position over time. **(A)** Control tadpole, **(B)** tadpole acutely exposed to 1 µM CPF for 1 hour, and **(C)** tadpole chronically exposed to 1 µM CPF from stage 42 to stage 49. Control and acutely exposed tadpoles show sustained looping around the perimeter of the well, whereas the chronically exposed tadpole shows irregular, disorganized movement consistent with seizure-like activity. **(D)** Total distance moved over 10 minutes for control, acute, and chronic CPF conditions. Acute CPF exposure significantly increased total distance traveled compared to controls, whereas chronic exposure did not differ from controls. **(E)** Fraction of active time spent performing each behavior type — looping (circling the perimeter), seizure-like (rapid, irregular swimming with frequent C-bends), and other (darting, spiraling, and undefined movements combined) — for control, acute CPF, and chronic CPF conditions. Chronically exposed tadpoles spent significantly less time looping and significantly more time in seizure-like activity compared to both controls and acutely exposed tadpoles. Bars represent mean ± SEM. ** p < 0.01, *** p < 0.001.

First, we investigated whether CPF exposure affected swimming distance. Measuring the total distance swum over 10 minutes, we found that tadpoles acutely exposed to CPF moved a greater average distance compared to controls (Fig. 2D; controls: 330.7 ± 205.6 cm (n=42); acute: 521.1 ± 150.6 cm (n=12), p= 0.0062). In contrast, tadpoles chronically exposed to CPF during development no longer showed any differences in total distance as controls (Fig. 2D; chronic: 290.5 ± 139.8 cm (n=18), p = 0.7168), consistent with an acute effect of blocking acetylcholinesterase activity which could lead to increased levels of activity.

When active, tadpoles predominantly swim around the edge of their arena, occasionally darting across (Hänzi & Straka, 2018). We quantified the time each tadpole spent performing distinct behaviors during spontaneous swimming under different conditions. Using an automated method, we classified tadpole movements into looping (circling the perimeter), darting (crossing the well), seizure-like (rapid, irregular swimming with many C-bends;(Thompson & Aizenman, 2024)), spiraling (slow circling mostly in the center), or undefined. We then calculated the fraction of active time spent on each behavior. Consistent with prior observations, control tadpoles predominantly performed looping behavior when active (Fig. 2E Fraction of active time: 0.82 ± 0.22). Tadpoles acutely exposed to CPF for 1 hour also predominantly performed looping behavior (Fraction of active time: 0.95 ± 0.06; p = 0.0762). In contrast, tadpoles chronically exposed to CPF spent less time looping compared to control and acutely exposed tadpoles (Fraction of active time: 0.37 ± 0.37; p < 0.0001). This decrease in looping behavior was associated with an increase in seizure-like behavior (Fig 2E; 0.42 ± 0.39) compared to both controls (0.05 ± 0.06; p < 0.0001) and acutely exposed tadpoles (0.01 ± 0.02; p < 0.0001). There was no significant change in the fraction of time spent on other behaviors between controls and CPF-exposed tadpoles (p < 0.3263). These data suggest that acute CPF exposure increases tadpole swimming without changing behavior types, while chronic exposure profoundly affects behavior, inducing spontaneous seizure activity. The breakdown of time spent doing each different behavior across individual tadpoles is shown in **Fig. 3**, indicating that the effects of the different treatments were mostly consistent among individual animals.

**Figure 3.**
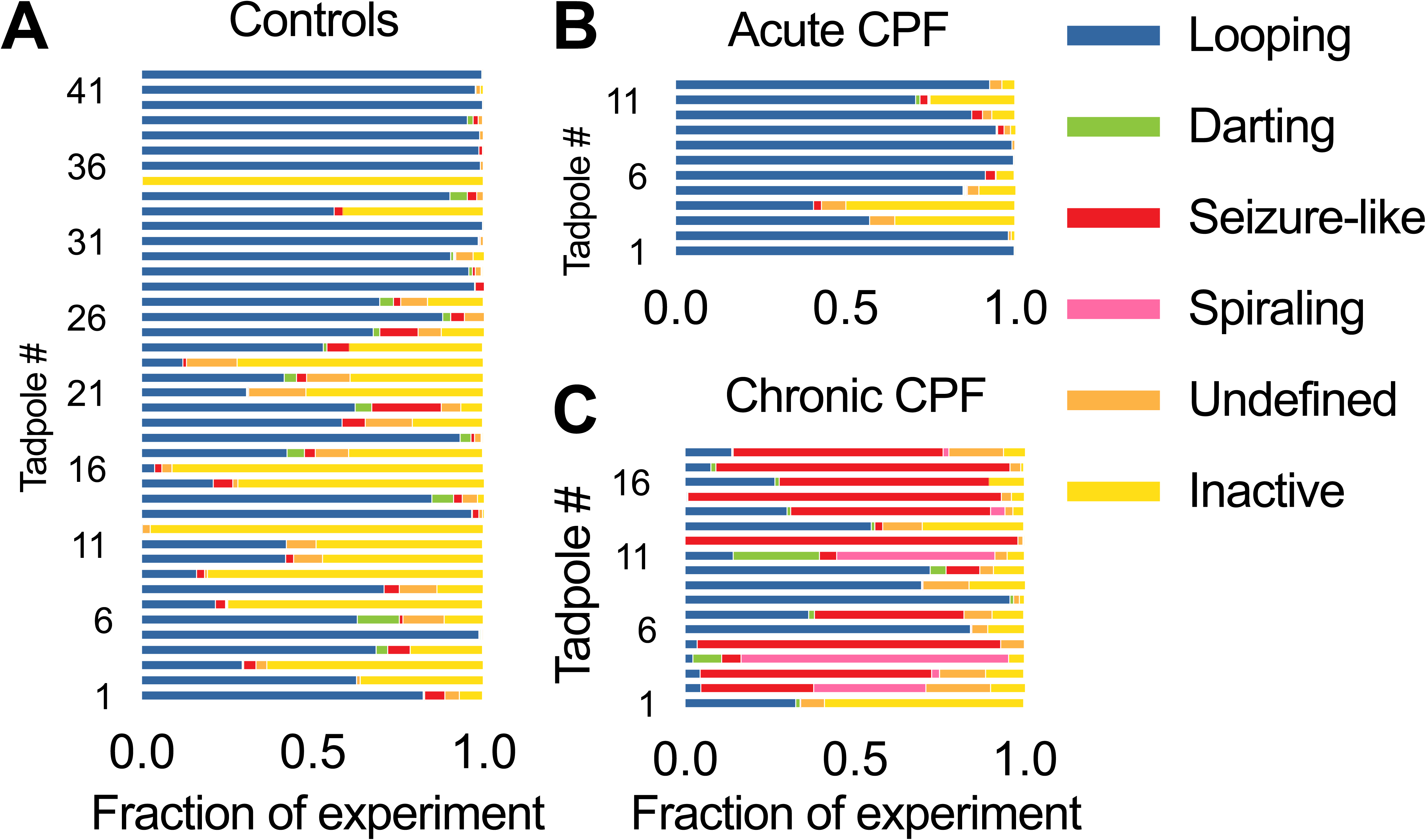
Behavioral composition is consistent across individual tadpoles within each condition. Stacked bar plots showing the fraction of the 10-minute recording each individual tadpole spent in each behavior category: looping, darting, seizure-like, spiraling, undefined, and inactive. **(A)** Control tadpoles, **(B)** tadpoles acutely exposed to 1 µM CPF for 1 hour, and **(C)** tadpoles chronically exposed to 1 µM CPF from stage 42 to stage 49. Each row represents one tadpole. Control and acutely exposed tadpoles predominantly performed looping behavior, whereas chronically exposed tadpoles showed a marked shift toward seizure-like activity. Variability within each group was modest, indicating that the behavioral effects of CPF exposure were consistent across individual animals.

Taken together, these data indicate that long-term developmental exposure to CPF results in a possible miswiring of neuronal circuitry that could lead, over time, to spontaneous seizures and abnormal schooling. Since these two behaviors are known to involve neural circuits in the optic tectum (James et al. 2015) we next performed electrophysiological recordings from tectal neurons in the tectum to see if we could find any differences.

### Single-cell electrophysiology

To examine differences in synaptic properties and intrinsic excitability in tectal neurons, we used an ex vivo whole brain preparation (Pratt & Aizenman, 2007). Brains were dissected either following a 1-hour CPF pre-exposure or after chronic developmental exposure. CPF was not present in the recording media. Whole-cell voltage and current clamp recordings were made from deep-layer tectal neurons that receive convergent visual and mechanosensory input (Deeg et al., 2009). Spontaneous excitatory and inhibitory postsynaptic currents (sEPSC, sIPSC) were collected at the GABA receptor reversal potential (-45mV) and the glutamate receptor reversal potential (+5 mV,(Wu et al., 1996)). sEPSC, sIPSC frequency, amplitude, and synaptic drive (frequency * amplitude) were calculated for acute 1-hour and chronic developmental CPF exposure.

While no differences were seen after acute exposure in excitatory transmission (Fig. 4A, C; Control, n=27 vs 1h CPF, n=13; sEPSC Amplitude: 6.1±0.3 vs 5.9±0.4 pA, p=0.55 M-W; sEPSC frq: 6.3±1.2 vs 4.9±1.1 events/sec, p=0.71; synaptic drive: 32.5±5.9 vs 32.6±10.3, p=0.73 M-W), there was a significant decrease in the frequency and overall synaptic drive of inhibitory responses (Fig. 4B,C; Control, n=27 vs 1h CPF, n=13; sIPSC Amplitude: 8.3±0.6 vs 7.5±0.6 pA, p=0.34 M-W; sIPSC frq: 11±0.7 vs 8.4±0.5 events/sec, p=0.002; synaptic drive: 93.1±1.4 vs 62.5±6.7, p=0.009 M-W), indicating a change in the balance of excitation to inhibition (E/I). After chronic exposure, this relative difference in the E/I balance persisted, with no changes in excitatory transmission (Fig. 5A, C; Control, n=33 vs 1h CPF, n=17; sEPSC Amplitude: 5.5±0.3 vs 4.9±0.2 pA, p=0.12 M-W; sEPSC frq: 2.6±0.5 vs 1.6±0.4 events/sec, p=0.13; synaptic drive: 14.6±2.8 vs 8±2.4, p=0.08 M-W) but a decrease in the amount of inhibition (Fig. 5B, C Control, n=22 vs chronic CPF, n=17; sIPSC Amplitude: 6.2±0.5 vs 5.2±0.2 pA, p=0.22 M-W; sIPSC frq: 7.1±0.7 vs 5.1±0.7 events/sec, p=0.04; synaptic drive: 48.1±8 vs 26.1±3.6, p=0.039 M-W). Using current-clamp recordings, we also measured the intrinsic excitability of tectal neurons. Input/output curves were generated for tectal neurons by injecting depolarizing current steps (from 10-150 pA) from a -65 mV membrane potential and plotting the number of evoked action potentials (Fig. 6A, C). For each cell, we also calculated the maximum number of spikes each cell could generate (Fig 6B). Generally, stage 49 tectal neurons only generate 2-4 spikes when depolarized as they are still immature (Pratt & Aizenman, 2007). We found that after a 1-hour exposure to CPF, tectal neurons showed increased excitability as evidenced by the input/output curves (**p<0.0001 2-way ANOVA, multiple comparison results shown on figure** ) and an increase in the maximum number of spikes (**Control: 4±0.3 spikes, n=35; acute CPF 7.2±1 spikes, n=14; p=0.0018 MW**). However, after chronic exposure, this difference in excitability had disappeared showing no difference in the input/output curves (p=0.5 2-way ANOVA) or maximum number of spikes (**Control: 2.7±0.2 spikes, n=40; chronic CPF 3±0.3 spikes, n=40; p=0.86 MW**).

**Figure 4.**
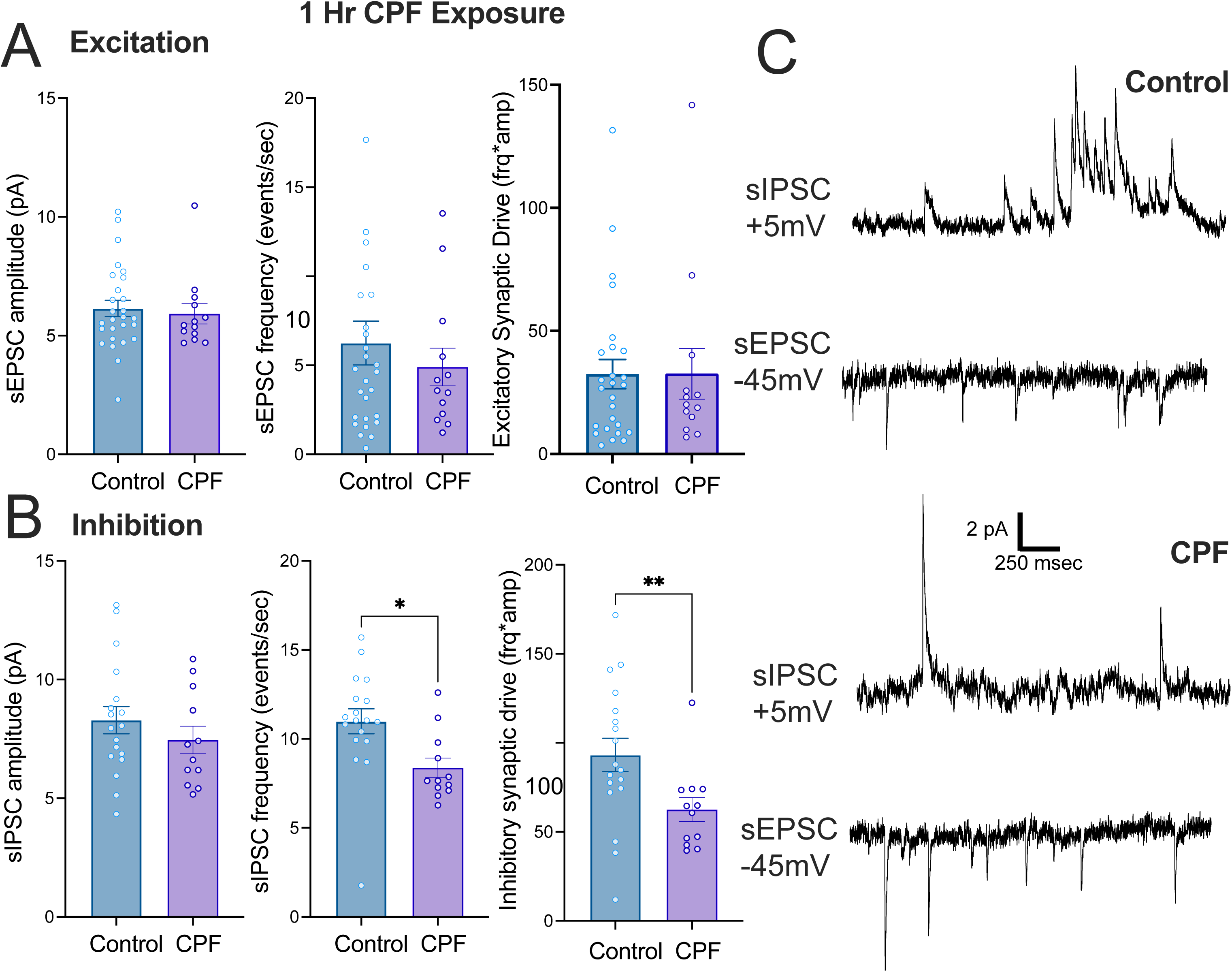
Acute CPF exposure selectively reduces inhibitory synaptic transmission in tectal neurons. Whole-cell voltage-clamp recordings of spontaneous excitatory and inhibitory postsynaptic currents (sEPSCs and sIPSCs) from deep-layer tectal neurons following 1-hour exposure to 1 µM CPF. sEPSCs were recorded at −45 mV and sIPSCs at +5 mV, corresponding to the reversal potentials for inhibitory and excitatory currents, respectively. **(A)** Excitatory transmission was unaffected by acute CPF exposure. sEPSC amplitude, frequency, and synaptic drive (frequency × amplitude) did not differ between control and CPF-exposed cells. **(B)** Inhibitory transmission was reduced by acute CPF exposure. sIPSC amplitude was unchanged, but sIPSC frequency and overall inhibitory synaptic drive were significantly decreased compared to controls. **(C)** Representative voltage-clamp traces of sIPSCs (recorded at +5 mV) and sEPSCs (recorded at −45 mV) from control and CPF-exposed tectal neurons. Bars represent mean ± SEM; circles represent individual cells. * p < 0.05, ** p < 0.01.

**Figure 5.**
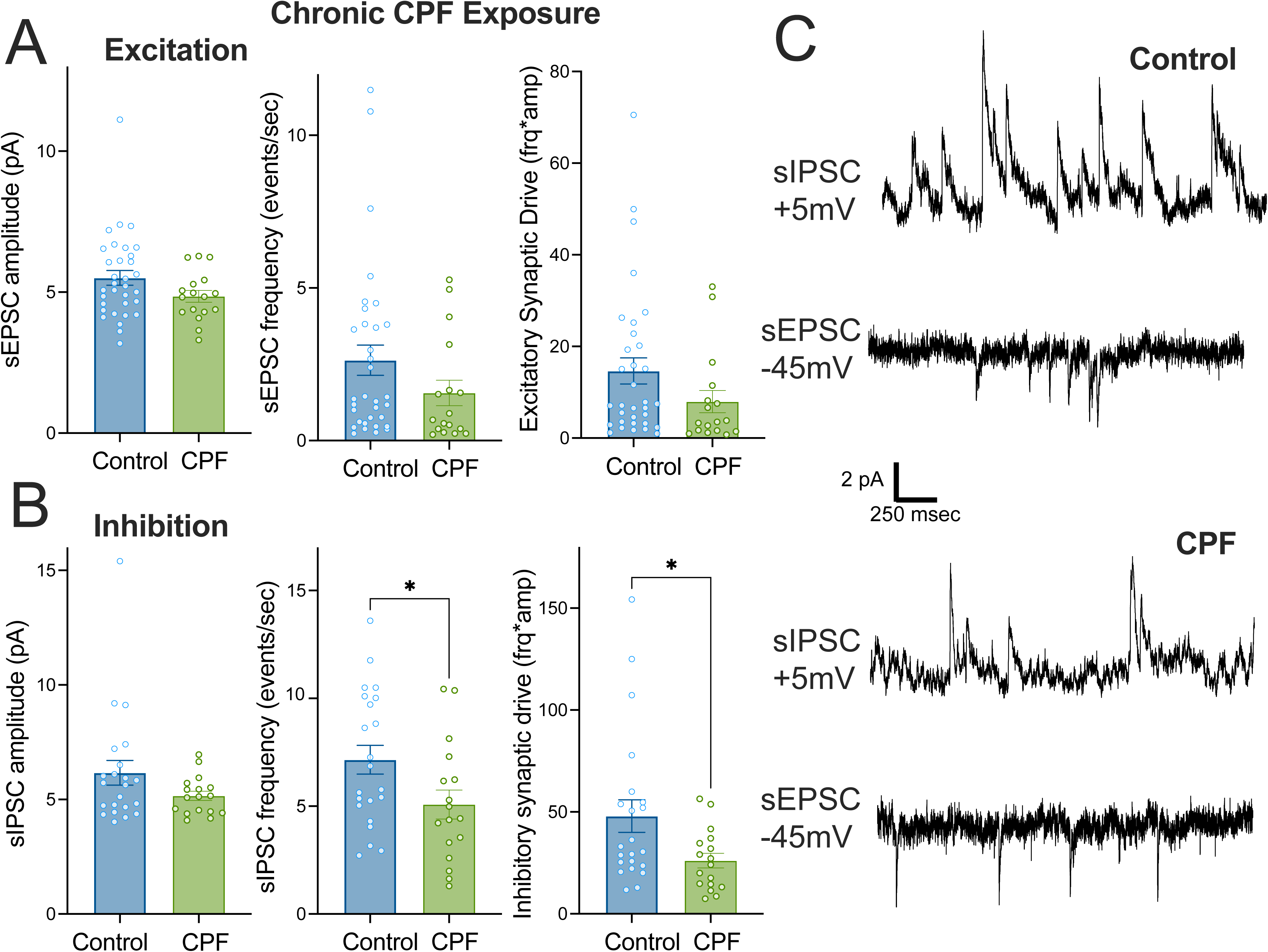
Chronic developmental CPF exposure produces a persistent reduction in inhibitory synaptic transmission. Whole-cell voltage-clamp recordings of spontaneous excitatory and inhibitory postsynaptic currents (sEPSCs and sIPSCs) from deep-layer tectal neurons following chronic CPF exposure (1 µM, stage 42 to stage 49). sEPSCs were recorded at −45 mV and sIPSCs at +5 mV. **(A)** Excitatory transmission was unaffected by chronic CPF exposure. sEPSC amplitude, frequency, and synaptic drive (frequency × amplitude) did not differ significantly between control and CPF-exposed cells. **(B)** Inhibitory transmission remained reduced following chronic exposure. sIPSC amplitude was unchanged, while sIPSC frequency and overall inhibitory synaptic drive were significantly decreased compared to controls. **(C)** Representative voltage-clamp traces of sIPSCs (recorded at +5 mV) and sEPSCs (recorded at −45 mV) from control and CPF-exposed tectal neurons. Bars represent mean ± SEM; circles represent individual cells. * p < 0.05.

**Figure 6.**
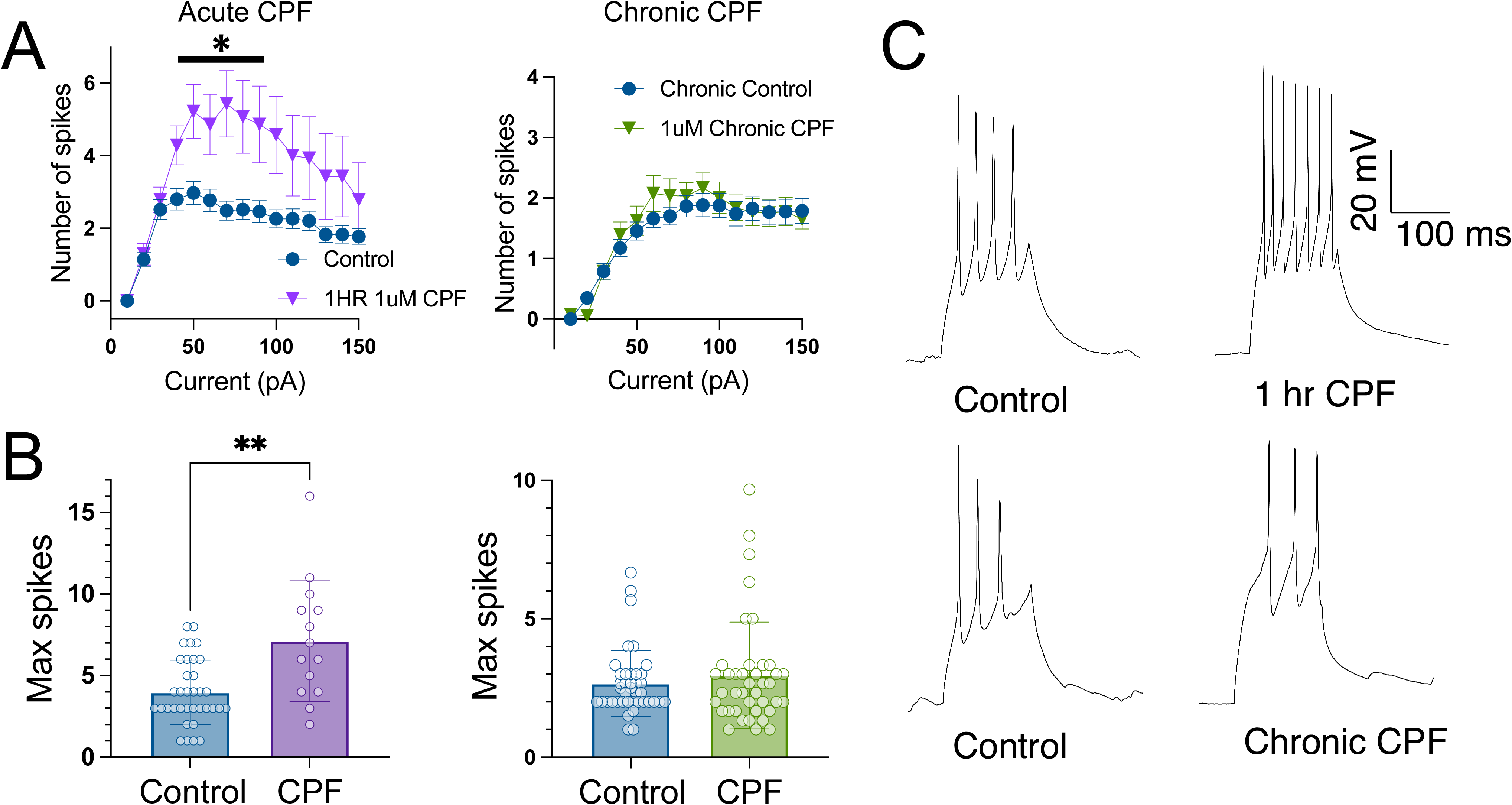
Acute CPF exposure transiently increases intrinsic excitability of tectal neurons, but this effect is not maintained after chronic exposure. Whole-cell current-clamp recordings from deep-layer tectal neurons. Input/output curves were generated by injecting depolarizing current steps (10–150 pA) from a holding potential of −65 mV and counting the number of evoked action potentials. **(A)** Input/output curves for acute CPF exposure (left; control vs. 1-hour 1 µM CPF) and chronic CPF exposure (right; control vs. chronic 1 µM CPF, stage 42 to stage 49). Acute CPF exposure significantly increased the number of evoked spikes across a range of current injections, whereas chronic exposure showed no difference from controls. The horizontal bar in the left panel indicates the range of current injections at which the two groups differed significantly in post-hoc multiple comparisons. **(B)** Maximum number of spikes evoked per cell for acute CPF (left) and chronic CPF (right) conditions. Acutely exposed neurons fired significantly more maximum spikes than controls, while chronically exposed neurons did not differ from controls. **(C)** Representative voltage traces showing action potentials evoked by a 200 msec depolarizing current injection from control, acute CPF, chronic control, and chronic CPF neurons. Bars represent mean ± SEM; circles represent individual cells. Points on input/output curves represent mean ± SEM. ** p < 0.01.

Both changes in the E/I balance and increases in intrinsic excitability have been proposed to underlie the generation of seizures during epilepsy (Žiburkus et al., 2013). However, these alone are not sufficient to explain the development of seizures after developmental exposure to CPF, since tadpoles exposed to CPF for 1 hour do not exhibit seizure behavior, but they do show enhanced swimming activity, decreased inhibition and increased intrinsic excitability. It is possible, however, that a persistent alteration in the E/I balance leads to abnormal circuit formation during development, which in turn can underlie seizure generation (Gore et al., 2021). Furthermore, abnormal tectal circuit formation could also lead to abnormal multisensory processing, which is central to tectal function and tectally mediated behavior (Truszkowski et al., 2017). To test this possibility, we examined the dendritic architecture of tectal neurons after developmental exposure to CPF, as an index of neural circuit development.

### Dendritic architecture

We used sparse electroporation of GFP-expressing plasmids (see methods) to drive expression of GFP in individual tectal neurons in both control and CPF-reared tadpoles. Following the CPF exposure period, we used in vivo confocal microscopy to reconstruct their dendritic arbors and analyze their morphology (Fig. 7A). When we measured the total dendritic branch length (control, 228.7 ± 16.7, n = 18; CPF, 236.0 ± 25.6, n = 19; p = .816), the average branch length (control, 22.47 ± 2.15, n = 18; CPF, 21.32 ± 1.56, n = 19; p = .667), and the average number of branches per cell (control, 12.56 ± 2.34, n = 18; CPF, 11.47 ± 1.32, n = 19; p = .686), we found no significant differences between the two groups (Fig. 7B).

**Figure 7.**
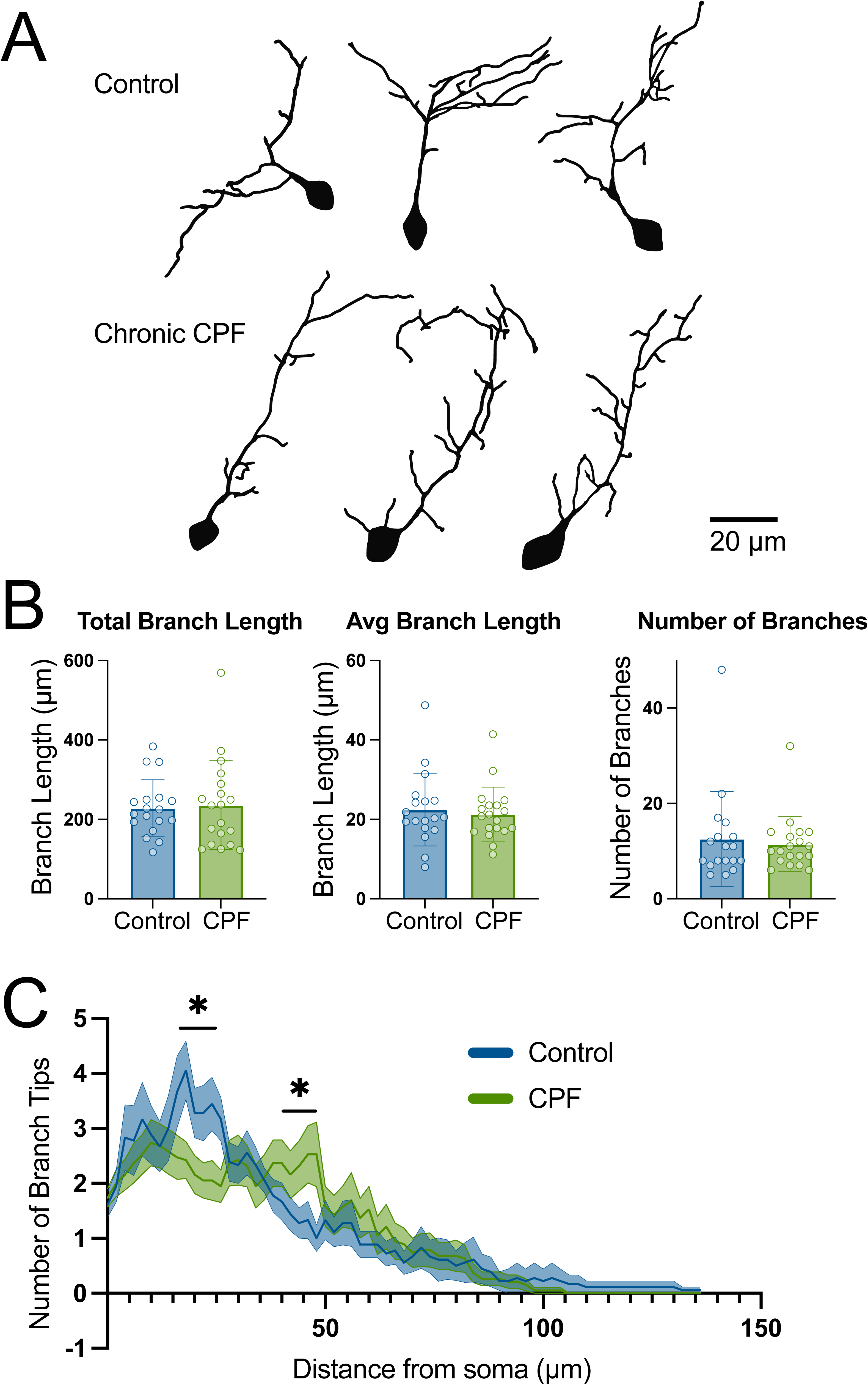
Chronic developmental CPF exposure alters the spatial distribution of dendritic branching without changing overall arbor size. Single tectal neurons were sparsely labeled by whole-brain electroporation with pCALNL-GFP and pCAG-Cre plasmids and imaged in vivo by confocal microscopy at stage 49. **(A)** Representative 3D reconstructions of GFP-expressing tectal neurons from control (top) and chronically CPF-exposed (bottom) tadpoles (1 µM CPF, stage 42 to stage 49). Scale bar, 20 µm. **(B)** Quantification of total dendritic branch length, average branch length, and number of dendritic branches per cell. None of these metrics differed significantly between control and CPF-exposed neurons (control, n = 18; CPF, n = 19). **(C)** Three-dimensional Sholl analysis showing the number of dendritic branch tips as a function of distance from the soma. Control neurons showed branch tips concentrated at a principal location proximal to the soma, whereas CPF-exposed neurons showed branch tips distributed more broadly along the primary dendrite. Lines represent mean and shaded regions represent SEM. Horizontal bars indicate ranges of distance from the soma at which the two groups differed significantly in post-hoc multiple comparisons (two-way ANOVA, p < 0.02). Bars in B represent mean ± SEM; circles represent individual cells. * p < 0.05.

However, a three-dimensional Sholl analysis of the branching patterns showed significant differences between the experimental groups (two-way ANOVA, p < .02, locations where distributions differ significantly are indicated in the figure). Specifically, control tadpoles tended to have dendritic branches concentrated in a principal location relative to the soma, whereas CPF-reared tadpoles exhibited branches dispersed throughout the primary dendrite (Fig. 7C,A). This difference in regional branching distribution suggests that CPF exposure leads to abnormal development of tectal dendrites, which is consistent with abnormal development of connectivity within the tectum (James et al., 2015). Such changes in tectal connectivity have been linked to the emergence of hyperexcitable circuits and increased seizure susceptibility.

## Discussion

Overall, we found that chronic developmental exposure to low concentrations of CPF resulted in multiple abnormal neurodevelopmental outcomes. Tadpoles chronically exposed to CPF exhibited significant alterations in swimming and schooling behaviors, including increased seizure-like activity accompanied by reduced exploratory looping. Developmental exposure also impaired schooling, as indicated by a failure to align with nearest neighbors.

Single-cell analyses of dendritic morphology suggest that neural circuitry is altered following developmental CPF exposure. Sholl analysis of dendritic branching patterns revealed significant differences in CPF-reared animals, with altered patterns of dendritic branch emergence from the primary shaft. These changes indicate abnormal circuit formation within the tectum, which could contribute to the observed behavioral alterations.

Finally, single-cell electrophysiological recordings revealed an acute increase in intrinsic excitability that was subsequently compensated for following chronic exposure. Despite this compensation, a persistent increase in network activity remained, as evidenced by a lasting decrease in inhibitory drive. This finding is consistent with a disruption of the excitation–inhibition balance and aligns with the observed seizure-like behavior. Taken together, these data suggest that developmental CPF exposure leads to miswiring of developing tectal circuits, resulting in an overall imbalance between excitatory and inhibitory synaptic drive. This circuit-level dysfunction is consistent with mechanisms proposed to underlie circuit dysfunction in ASD and other disorders (Bengoetxea de Tena et al., 2026). The acute increase in tectal neuron excitability and decreased inhibition also correlate well with the elevated swimming activity, but absence of seizures, seen after 1 hour exposure of CPF. This suggests the development of seizure activity reflects a developmental miswiring, as opposed to simply an increase in network excitability.

CPF is an organophosphate that exerts its main effects by inhibiting AChE (Lan et al., 2017; Silva, 2020). In the brain, ACh is an important neuromodulator. ACh exerts its diverse effects in the brain by acting through metabotropic and ionotropic receptors. Nicotinic acetylcholine receptors (nAChRs) are expressed within the optic tectum, particularly in retinorecipient layers involved in visual processing, where they are localized both on tectal neurons and presynaptically on retinal ganglion cell (RGC) terminals (Butt et al., 2000; Edwards & Cline, 1999a). Receptors containing α7 subunits are of particular interest because they are highly calcium permeable and mediate excitatory cholinergic signaling, potentially influencing synaptic plasticity and circuit refinement within the developing tectum. In addition to nAChRs, both excitatory and inhibitory muscarinic acetylcholine receptors (mAChRs) are distributed throughout the tectum. Inhibitory muscarinic receptors are broadly expressed, whereas other muscarinic receptor populations are enriched within retinorecipient layers, likely on tectal neurons themselves (Butt et al., 2001). Together, these receptor populations allow acetylcholine (ACh) to exert both excitatory and inhibitory neuromodulatory effects on tectal circuitry, thereby regulating visual processing and neural network activity during development.

ACh is an important regulator of visual circuit development in the tectum. Most cholinergic inputs to the tectum originate in the nucleus isthmi, another midbrain structure (Desan et al., 1987; Wallace et al., 1990). Release of ACh from nucleus isthmi projections enhances visual transmission in the tectum(Edwards & Cline, 1999b; Gruberg et al., 1991; King & Schmidt, 1991) This pathway is also critical for maintaining the topography of the retinotectal map, which is refined through activity-dependent mechanisms during development (Schmidt, 1985; Tu et al., 2000). ACh plays a similar role in the developing mammalian visual system, where it regulates visual circuit development and plasticity (Bear & Singer, 1986; Bröcher et al., 1992; Kirkwood et al., 1999). Consistent with these developmental functions, alterations in cholinergic transmission are known to disrupt normal neural circuit development (Heath et al., 2010).

Given the prominent role of ACh in regulating tectal excitability, visual transmission, and map refinement, excess ACh following CPF exposure could disrupt circuit function through both muscarinic and nicotinic receptor-dependent mechanisms. Some subtypes of metabotropic receptors have inhibitory effects, preventing further release of ACh and reducing glutamate at cortiocortical and cortiostriatal synapses. Other types have excitatory effects, increasing dopamine release at striatal synapses and increasing excitability of pyramidal cells. Nicotinic ACh receptors (nAChRs) also play a modulatory role in the central nervous system. ACh has also been implicated in several important processes related to neurodevelopment and synaptic plasticity. Pre-synaptic nAChRs may increase the release of several neurotransmitters, such as dopamine, serotonin, glutamate, and GABA, while post-synaptic ones can increase neuronal firing rate and facilitate LTP ( see (Picciotto et al., 2012)). Since nAChRs are permeable to Ca^2+^, this gives ACh the ability to facilitate neurotransmitter release on the pre-synaptic side by directly increasing calcium and indirectly through opening voltage-gated calcium channels. Whereas postsynaptic activation will trigger signalling cascades related to plasticity (McKay et al., 2007).

How might AChE inhibition produce the specific pattern of persistent GABAergic deficits we observed? One particularly compelling possibility involves the developmental switch in chloride transporter expression that underlies the maturation of GABAergic inhibition. Early in development, neurons express the chloride importer NKCC1, which maintains a high intracellular chloride concentration and causes GABA to act as a depolarizing neurotransmitter. The subsequent switch to KCC2 expression lowers intracellular chloride and renders GABA inhibitory (Z. Liu et al., 2007; Picciotto et al., 2012). nAChR signaling has been implicated in driving this transition, raising the possibility that excess cholinergic activation during CPF exposure could disrupt the normal timing of the NKCC1-to-KCC2 switch. Such a disruption would be expected to produce lasting alterations in GABAergic transmission, consistent with the persistent reduction in inhibitory synaptic drive we observed even after acute excitability changes had normalized. Testing this hypothesis directly, for example, by examining the developmental timing of chloride transporter expression in CPF-exposed tadpoles, represents an important direction for future work.

Although cholinergic mechanisms provide a plausible framework for many of our findings, CPF likely acts through additional pathways. Several studies have found CPF affects neurodevelopment even at doses below the threshold for AChE inhibition. For example, one study found that there was an increase in serotonin turnover in rats exposed early in development to CPF, suggesting an alteration in synaptic functioning (Aldridge et al., 2005). This study also found that there was a significant reduction in dopamine in the hippocampus in rats exposed to this low dose of CPF. Other research has observed deficits in locomotor activity, cognition, and learning and memory in rodents exposed to doses below the AChE threshold (Aldridge et al., 2005; Silva, 2020). This suggests that CPF may have other, off-target effects that impact the nervous system using a non-cholinergic mechanism. Altered serototonergic signalling has been shown to also affect development of tectal circuitry and is necessary for normal visually guided behavior (Bruno et al., 2022; K. Liu et al., 2021).

In sum, we show that developmental exposure to CPF results in an early change in neuronal excitability, which persists as an altered E/I balance and increased network excitability. This is accompanied by altered dendritic organization and manifests behaviorally as seizure-like activity and abnormal social behavior, both hallmarks of various neurodevelopmental disorders including autism. While our analysis focused on tectal circuits and tectally mediated behavior, this change likely also occurs across other brain regions exposed to CPF during development. These data illustrate how small alterations at the synaptic and intrinsic excitability levels can cascade into broad behavioral abnormalities, providing a plausible mechanistic explanation for how early developmental exposure to AChE-inhibiting pesticides can contribute to neurodevelopmental disorders. These findings have particular relevance for understanding the risks of occupational CPF exposure during pregnancy in agricultural settings and agricultural communities, and for ongoing debates about the regulation of organophosphate pesticides.

## Acknowledgements

This study was completed during the height of the COVID pandemic, with funding from Brown University.

## Author Contributions

VLIII: electrophysiology, behavior, experimental design, data analysis, manuscript preparation. AR: dendritic imaging, experimental design, data analysis, manuscript preparation. AT: analysis of swimming behavior, manuscript preparation. ZP: manuscript preparation, data analysis, CA: experimental design, data analysis, manuscript preparation.

## Notes

### Competing Interest Statement

The authors have declared no competing interest.

